# Differential gene expression, including *Sjfs800,* in *Schistosoma japonicum* females before, during, and after male-female pairing

**DOI:** 10.1101/452458

**Authors:** Fengchun Liu, Han Ding, Jiaming Tian, Congyu Zhou, Fei Yang, Wei Shao, Yinan Du, Xin Hou, Cuiping Ren, Jijia Shen, Miao Liu

## Abstract

Schistosomiasis is a prevalent but neglected tropical disease caused by parasitic trematodes of the genus *Schistosoma,* with the primary disease-causing species being *S. haematobium*, *S. mansoni,* and *S. japonicum.* Male-female pairing of schistosomes is necessary for sexual maturity and the production of a large number of eggs, which are primarily responsible for schistosomiasis dissemination and pathology. Here, we used microarray hybridization, bioinformatics, quantitative PCR, in situ hybridization, and gene silencing assays to identify genes that play critical roles in *S. japonicum* reproduction biology, particularly in vitellarium development, a process that affects male-female pairing, sexual maturation, and subsequent egg production. Microarray hybridization analyses generated a comprehensive set of genes differentially transcribed before and after male-female pairing. Although the transcript profiles of females were similar 16 and 18 days after host infection, marked gene expression changes were observed at 24 days. The 30 most abundantly transcribed genes on day 24 included those associated with vitellarium development. Among these, genes for female-specific 800 *(fs800*), *eggshell precursor protein*, and superoxide dismutase *(cu-zn-SOD)* were substantially upregulated. Our in situ hybridization results in female *S. japonicum* indicated that *cu-zn-SOD* mRNA was highest in the ovary and vitellarium, *eggshell precursor protein* mRNA was expressed in the ovary, ootype, and vitellarium, and *Sjfs800* mRNA was observed only in the vitellarium, localized in mature vitelline cells. Knocking down the *Sjfs800* gene in female *S. japonicum* by approximately 60% reduced the number of mature vitelline cells, decreased rates of pairing and oviposition, and decreased the number of eggs produced in each male-female pairing by about 50%. These results indicate that *Sjfs800* is essential for vitellarium development and egg production in *S. japonicum* and suggest that *Sjfs800* regulation may provide a novel approach for the prevention or treatment of schistosomiasis.

**Author Summary:** Schistosomiasis is a common but largely unstudied tropical disease caused by parasitic trematodes of the genus *Schistosoma.* The eggs of schistosomes are responsible for schistosomiasis transmission and pathology, and the production of these eggs is dependent on the pairing of females and males. In this study, we determined which genes in *Schistosoma japonicum* females were differentially expressed before and after pairing with males, identifying the 30 most abundantly expressed of these genes. Among these 30 genes, we further characterized those in female *S. japonicum* that were upregulated after pairing and that were related to reproduction and vitellarium development, a process that affects male-female pairing, sexual maturation, and subsequent egg production. We identified three such genes, *S. japonicum* female-specific 800 *(Sjfs800), eggshell precursor protein,* and superoxide dismutase, and confirmed that the mRNAs for these genes were primarily localized in reproductive structures. By using gene silencing techniques to reduce the amount of *Sjfs800* mRNA in females by about 60%, we determined that *Sjfs800* plays a key role in development of the vitellarium and egg production. This finding suggests that regulation of *Sjfs800* may provide a novel approach to reduce egg counts and thus aid in the prevention or treatment of schistosomiasis.

## Introduction

Schistosomiasis, also known as bilharzia, is a tropical disease caused by parasitic trematodes of the genus *Schistosoma.* Although it is one of the most prevalent tropical infectious diseases, with more than 240 million people in 78 countries infected and approximately 800 million people at risk, schistosomiasis has been drastically understudied [1-3]. The primary disease-causing species of *Schistosoma* are *S. haematobium*, *S. mansoni,* and *S. japonicum*, the latter of which is distributed in China, Indonesia, and the Philippines [1-3]. Disease burden assessments for schistosomiasis, based on the extent of end-organ damage and the associated morbidities related to malnutrition and chronic inflammation, indicate that the annual number of disability-adjusted life years lost is approximately 70 million [4]. Current control of schistosomiasis depends largely on a single drug, praziquantel; however, reliance on a single drug produces a precarious situation. Indeed, some studies have shown that isolates of schistosomes have reduced susceptibility to praziquantel [5-7]. Thus, additional novel strategies are urgently needed to prevent and control schistosomiasis.

*S. japonicum* has a complex developmental cycle that involves an aquatic snail as an intermediate host and a mammalian definitive host. In contrast to other trematode species, these parasites are unique in that males and females need to pair to continue development. Pairing of schistosome females and males promotes female reproductive system maturation and the production of eggs, which are a primary means of schistosomiasis transmission and immunopathological lesions [8-10]. Maturation and maintenance of normal reproductive function in female *S. mansoni* require permanent pairing with the male. During pairing, germ cells in the reproductive organ differentiate into oocytes or vitellocytes, and some chemical or tactile stimulus exchange occurs between the male and female, leading to a cascade of changes during the pairing process [11-16]. However, the effects on female reproductive system development and the molecular mechanisms underpinning male-female pairing have not been completely determined, leaving myriad questions that require further study.

Ongoing work in our laboratory has indicated that during the development of *S. japonicum,* no male-female pairing occurs up to 16 days after the host is infected. Some pairing occurs 17 days post infection (dpi), and pairing is common 18 dpi. Paired females begin laying eggs approximately 24 dpi. Therefore, in the present study, to identify genes that likely contribute to pairing and reproduction, we used microarray technology to determine differential gene expression in females 16, 18, and 24 dpi. We identified genes that play critical roles in the development of the vitellarium and in the production of eggs, providing a clearer understanding of gene regulation before and after male-female pairing in the *S. japonicum* female and insights on schistosome reproduction biology.

## Materials and methods

### Ethical statement

All procedures performed on animals within this study were conducted using animal husbandry guidelines approved by the Animal Ethics Committee of Anhui Medical University (LLSC20140060).

### Animals and parasites

Freshly shed wild-type cercariae of *S. japonicum* were harvested from infected *Oncomelania hupensis* that were purchased from the Hunan Institute of Parasitic Diseases in Yueyang, China. Female Kunming mice (6-8 weeks old) and New Zealand rabbits (4 months old) were obtained from the Laboratory Animal Center of Anhui Medical University. New Zealand rabbits and female Kunming mice were infected with 1000 or 50 cercariae, respectively, via the skin of the abdomen. After 16, 18, 24, 28, or 42 dpi the worms were washed out the hepatic portal vein using perfusion techniques. Male and female worms were manually separated. In order to collect *S. japonicum* eggs, liver tissues from rabbits 6 weeks post infection were homogenized and then subjected to consecutive fractional filtration. The filtrate was centrifuged. The supernatant and the tissue-containing layers were removed, leaving the egg-containing layer, which was diluted in 1.2% saline and passed through a nylon net (300 mesh, i.e., 300 holes per inch). All parasite samples were soaked in RNAlater (Invitrogen, Thermo Scientific, USA) and stored at −80°C until they were used for total RNA extraction.

### RNA extraction, amplification, and labeling

Total RNA was extracted and purified using RNeasy micro kit (QIAGEN, #74004, GmBH, Germany) following the manufacturer’s instructions, and the overall RNA quality was assessed using denaturing gel electrophoresis (Agilent Technologies, Santa Clara, CA, USA). Total RNA was amplified and labeled using a Low Input Quick Amp Labeling Kit, one-color (Agilent Technologies, #5190-2305), following the manufacturer’s instructions. Labeled cRNA was purified using a RNeasy Mini Kit (QIAGEN, #74106).

### Microarray construction and hybridization and subsequent data analysis

A schistosome genome-wide microarray was used for profiling gene expression in *S. japonicum* female 16, 18, and 24 dpi. Microarrays were printed on the Agilent custom Schistosoma 4 x 44K chip (design ID: 048766). There were eight specimens; thus, eight chips in total were needed. Each slide was hybridized with 1.65 μg of Cy3-labeled cRNA using a Gene Expression Hybridization Kit (Agilent Technologies, #5188-5242) with a Hybridization Oven (Agilent Technologies, #G2545A) according to the manufacturer’s instructions. After 17 hours of hybridization had elapsed, the slides were washed in staining dishes (Thermo Shandon, #121, Waltham, MA, USA) with a Gene Expression Wash Buffer Kit (Agilent Technologies, #5188-5327), following the manufacturers’ instructions. All of these aforementioned procedures were performed by Biotechnology Shanghai China. The slides were scanned with an Agilent Microarray Scanner (Agilent Technologies, #G2565CA) using the following default settings: dye channel, green; scan resolution, 5 μm; PhotoMultiplier Tube, 100% and 10%, 16 bit. Data were extracted with Feature ’ Extraction software, version 10.7 (Agilent Technologies). Raw data were normalized using the Quantile algorithm, GeneSpring software, version 11.0 (Agilent Technologies). Outlier probes were identified, and their contribution was reduced at the reported gene expression level. The expression value of a gene was a weighted average of all forward or reverse probe sets when both background correction and quantile normalization were performed.

### Bioinformatics analysis

The mRNA and expressed sequence tag transcripts highly enriched in *S. japonicum* 16, 18, and 24 dpi were retrieved from the National Center for Biotechnology Information Entrez Gene database (http://www.ncbi.nlm.nih.gov/gene)’ based on fold change (FC ≥ Signal A/Signal B) values (FC ≥2, three biological replicates; FC ≥3, two biological replicates). Student’s t-test was used to determine ; genes differentially expressed between one stage and the other *(p* < 0.05). All analyses were conducted using the online SBC Analysis System of Shanghai i Biotechnology Corporation (http://sas.shbio.com).

### Quantitative PCR (qPCR)

Thirteen genes whose expression levels were increased and two genes whose expression levels were decreased in females after pairing relative to those levels before pairing were selected for validation using qPCR. Total RNA (500 ng) from the females was reverse transcribed into first-strand cDNA using a PrimeScript RT ’ Reagent Kit (TaKaRa & Clontech, #RR037A, Japan) according to the manufacturer’s instructions. Each 10 μL of PCR reaction contained 10 μL of 2* SYBR Premix Ex Taq II, 1 μL of cDNA, 1.6 μL of the forward and reverse primer pair, 0.4 μL of 50* ROX Reference Dye, and 7 μL of sterile water. The PCR cycling conditions were as follows: 95°C for 30 s, followed by 40 cycles of 30 s denaturation at 95°C and 1 min annealing and extension at 60°C. A dissociation step (95°C for 15 s, 60°C for 1 min, and 95°C for 15 s) was performed to confirm the amplification specificity for each gene. A reliable reference gene for transcriptomic analysis of *S. japonicum,* proteasome 26S subunit, non-ATPase 4 *(PSMD4),* was used as a control gene in the assays. The PCR primers were designed using Primer Premier 5 software (Table 1). PCR reactions were performed in technical triplicates using the StepOnePlus Real-Time PCR System (Applied Biosystems). The relative expression level of each gene was analyzed using SDS 1.4 software (Applied Biosystems).

**Table 1.**
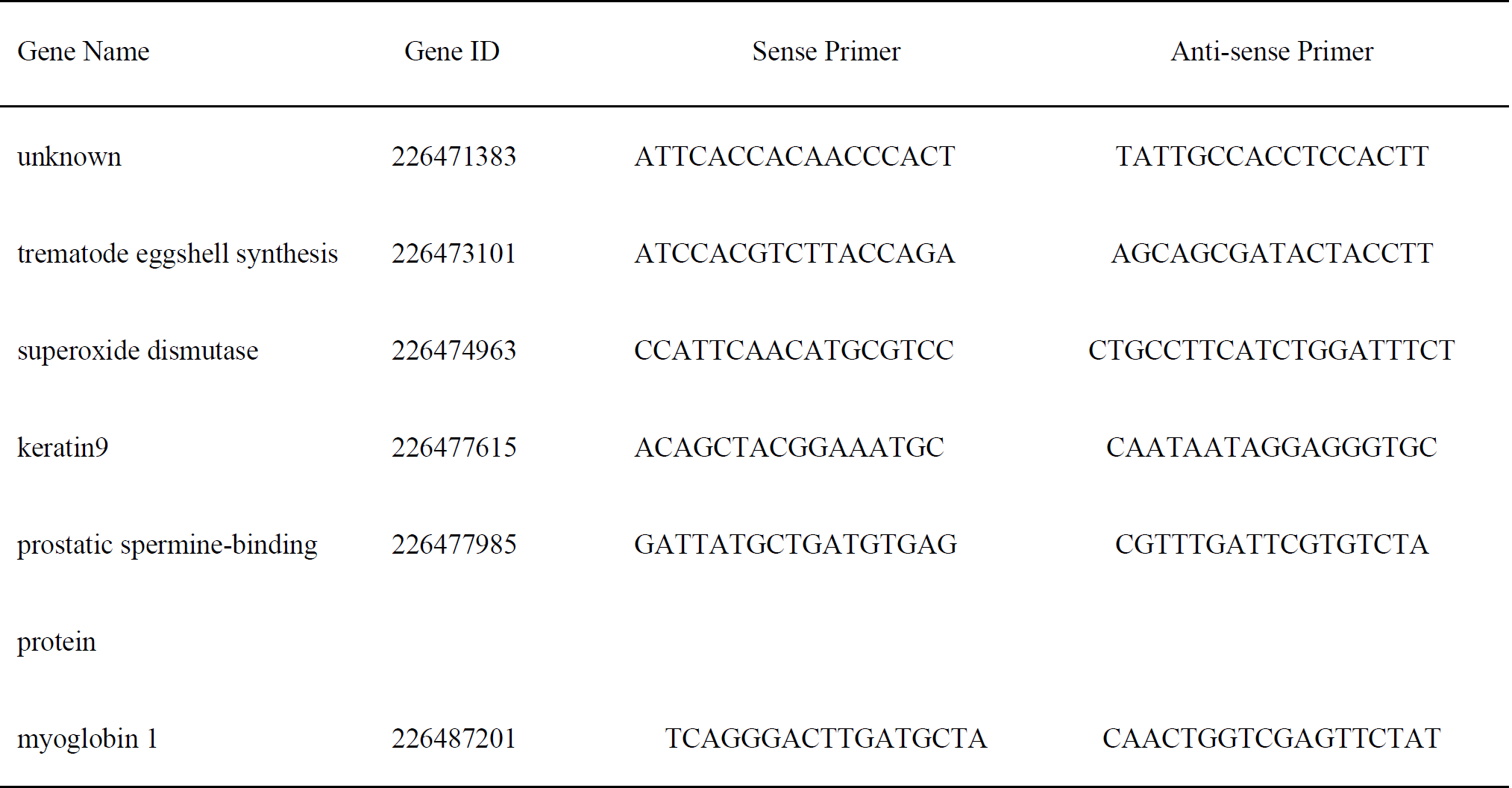
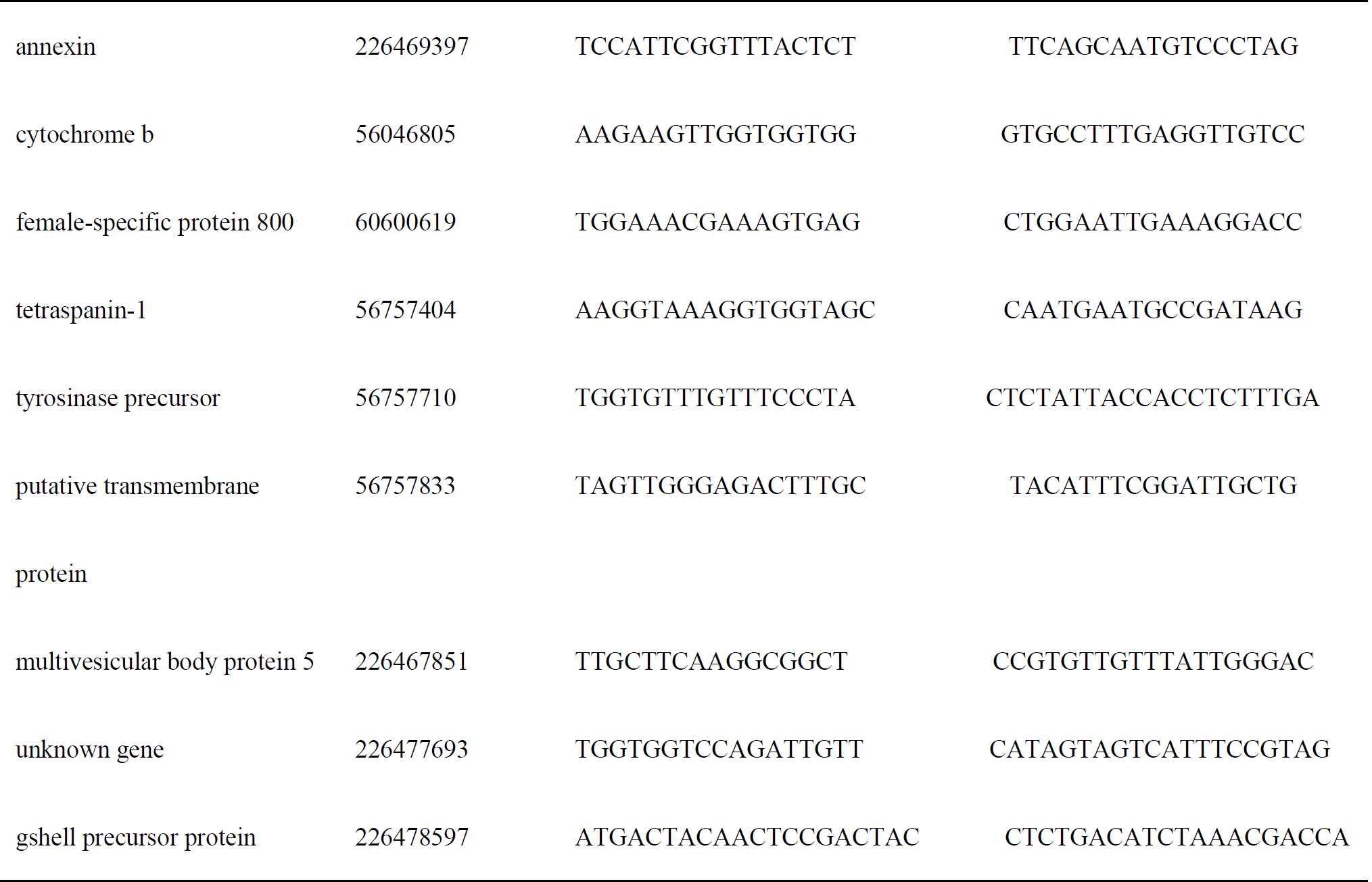
Quantitative PCR Primers (5’ to 3’).

Total RNAsfrom eggs, cercariae, schistosomula, and females 24 and 42 dpi were extracted using TRIzol reagent (Invitrogen) following the manufacturer’s instructions. The total RNA concentration and purity were measured using a NanoDrop 2000 (Thermo Fisher). Quantitative PCR was performed as described above using primer (Table 1) combinations to amplify gene transcripts of *S. japonicum* female-specific 800 *(fs800*), superoxide dismutase (*cu-zn-SOD*), and *eggshell precursor protein.*

### In situ hybridization

Riboprobes were synthesized according to previously published methods [17]. Briefly, probes were synthesized from restriction enzyme–digested DNA according to the orientation of the insert in pSPT18 using a DIG RNA Labeling Kit (SP6/T7) (Roche, #111750251910, Germany) labeled with digoxigenin. For whole-mount in situ hybridization, trematodes were fixed in 4% paraformaldehyde for 45 min and dehydrated in methanol. Following being bleached in 6% hydrogen peroxide in methanol to prevent tanning of the vitellaria, trematodes were permeabilized using proteinase k (TaKaRa & Clontech, #9034), incubated with prehybridization buffer (50% deionized formamide, 5× saline sodium citrate, 1 mg/mL yeast RNA, 1% Tween 20) for 2 h and then hybridized (prehybridization buffer with 10% dextran sulfate) with a riboprobe at 56°C for 20 h. Excess riboprobe was removed by washing in 2× and 0.2* saline sodium citrate, followed by blocking in blocking reagent (Roche, #11175041910). The bound riboprobe was detected after incubation of trematodes in antidigoxigenin alkaline phosphatase-conjugated antibody (Roche) diluted 1:5000 in blocking reagent overnight at 4°C. The unbound antibody was removed by washing in maleic acid buffer (100 mM maleic acid, 150 mM NaCl, and 0.1% Tween 20 at pH 7.5) for 4 h in twelve changes of buffer. After being washed, specimens were incubated in detection buffer (0.1 M Tris-HCl and 0.1 M NaCl at pH 9.5). Hybridization signals were detected by adding 200 μL of NBT/BCIP in detection buffer. After development, trematodes were washed in PBS then de-stained with 100% ethanol. Trematodes were mounted in 80% glycerol, and then microscopy and digital image capture were performed using an Olympus DP73 microscope.

### RNA interference

The siRNAs (21 base pairs) were designed using the *Sjfs800* mRNA sequence (GenBank accession No., FN313803.1) with the Thermo Fisher website software (https://rnaidesigner.thermofisher.com) and chemically synthesized by Shanghai GenePharma Co., Ltd. (China). Three siRNA sequences (siRNA1-3) that shared no homology with any other *S. japonicum* gene based on an online analysis with BLAST (National Center for Biotechnology Information) and the scrambled siRNA sequence are shown in Supplementary Table 1. The results of RNAi were tested using qPCR as previously described [18]. Briefly, Kunming mice were challenged percutaneously with 60-70 cercariae and were humanely killed 28 dpi. The worms were obtained by portal perfusion using RPMI 1640 medium at 37°C and were then incubated in 24-well plates (15 pairs/well) containing 1 mL of complete Basch medium, with half of the medium exchanged every day. The medium was supplemented with 10 KU/mL penicillin, 10 mg/ml streptomycin, 250 μg/mL amphotericin B (Sangon Biotech, #B13540732, China), and 10% fetal bovine serum (Gibco, #10091148, Thermo Scientific, USA). Lipofectamine RNAiMAX Transfection Reagent (Invitrogen) was used to transfect worms with one of the three *Sjfs800*-specific siRNAs, at a final concentration of 100 nM, or RNAase-free water (mock, no siRNA). The gene-silencing effect of each siRNA was determined using qPCR at the end of a 72-h cultivation period. The worms were then transfected with either the siRNA that was found to be most efficient or with the scrambled siRNA (control). The transfected worms were cultivated for up to 10 days and transfected again with siRNA1 on the fourth and seventh days; half of the medium was exchanged every day. After 10 days, all the eggs in the medium were collected and counted using light microscopy. Adult male-female pairings were also observed and counted. The siRNA silencing effects and morphological changes were measured using qPCR and confocal laser scanning microscopy (Leica TCS SP5, Germany), respectively.

### Confocal laser scanning microscopy

The confocal laser scanning microscopy procedure has been described previously [19]. After 10 days of *Sjfs800-specific* siRNA1 treatment, the male-female paired couples were separated manually and fixed separately in a solution of 95% alcohol, 3% formaldehyde, and 2% glacial acetic acid. The trematodes were then stained in 2.5% hydrochloric carmine (Ourchem, #71009382, China) for 16 h at 37°C and de-stained in 70% acidic ethanol. After dehydration in an ethanol series for 1 min in each concentration, parasites were cleared for 1 min each in 50% xylene diluted in ethanol and 100% xylene, and then whole-mounted with neutral balsam (Sinopharm Chemical #36645, China) on glass slides. The reproductive organ morphology of the trematodes was examined using confocal laser scanning microscopy, with a 470-nm longpass filter and a 488-nm He/Ne laser under reflection mode.

### Statistical analysis

Results were acquired from triplicate values representing three independent experiments with identical conditions. One-way analysis of variance and independent-sample *t* tests were used for data analysis with the SPSS, version 17.0, statistics software package. Data are expressed as the mean ± SEM, and *p* < 0.05 or *p* < 0.01 were deemed statistically significant.

## Results

### Microarray screening of differentially expressed genes in female *S. japonicum* before, during, and after pairing with males

The results of microarray analyses, the most comprehensive and informative probe assay design to date, indicated that after removing the duplicates, signal intensities were upregulated (FC ≥2), for 132 sequences during the pairing stage and for 198 sequences in paired female trematodes. Many mRNA transcripts were differentially expressed before, during and after pairing, with most of these genes elevated in expression after pairing. The most highly increased gene products after pairing were associated primarily with oxygen metabolism, the metabolic machinery of egg production, and vitellarium development. The 30 differentially expressed genes with highest expression levels after pairing are given in Table 2.

**Table 2.**
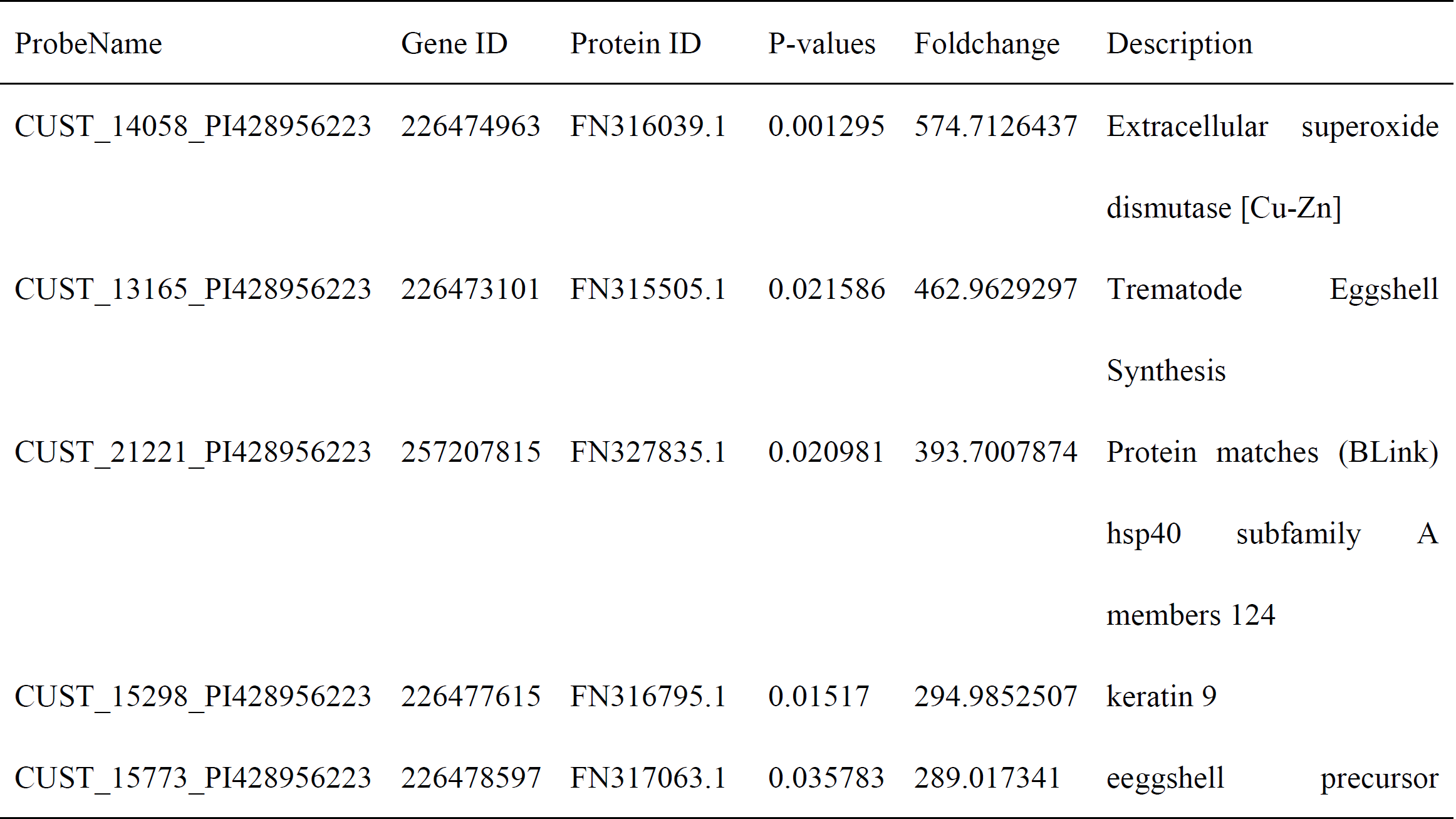
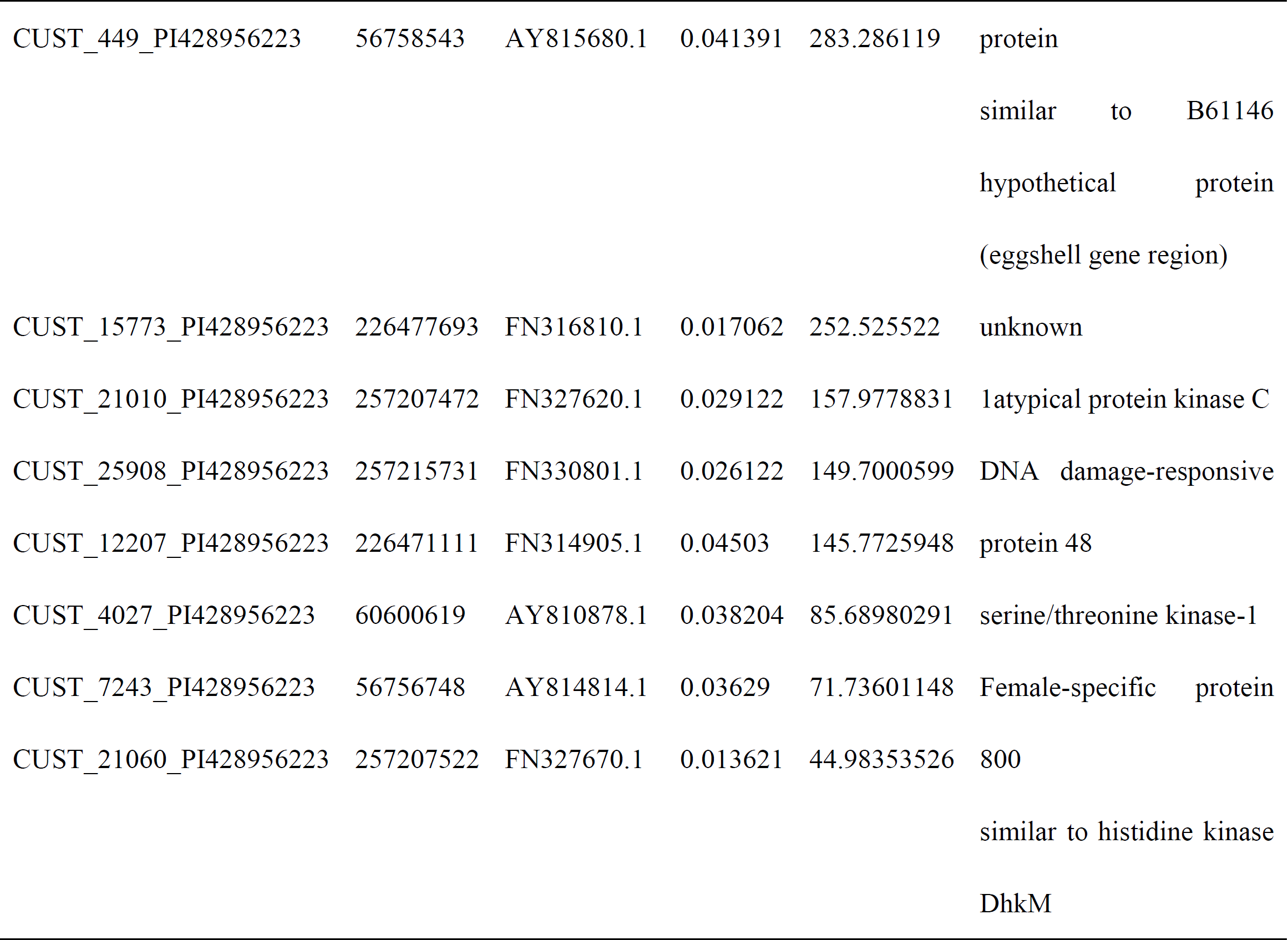
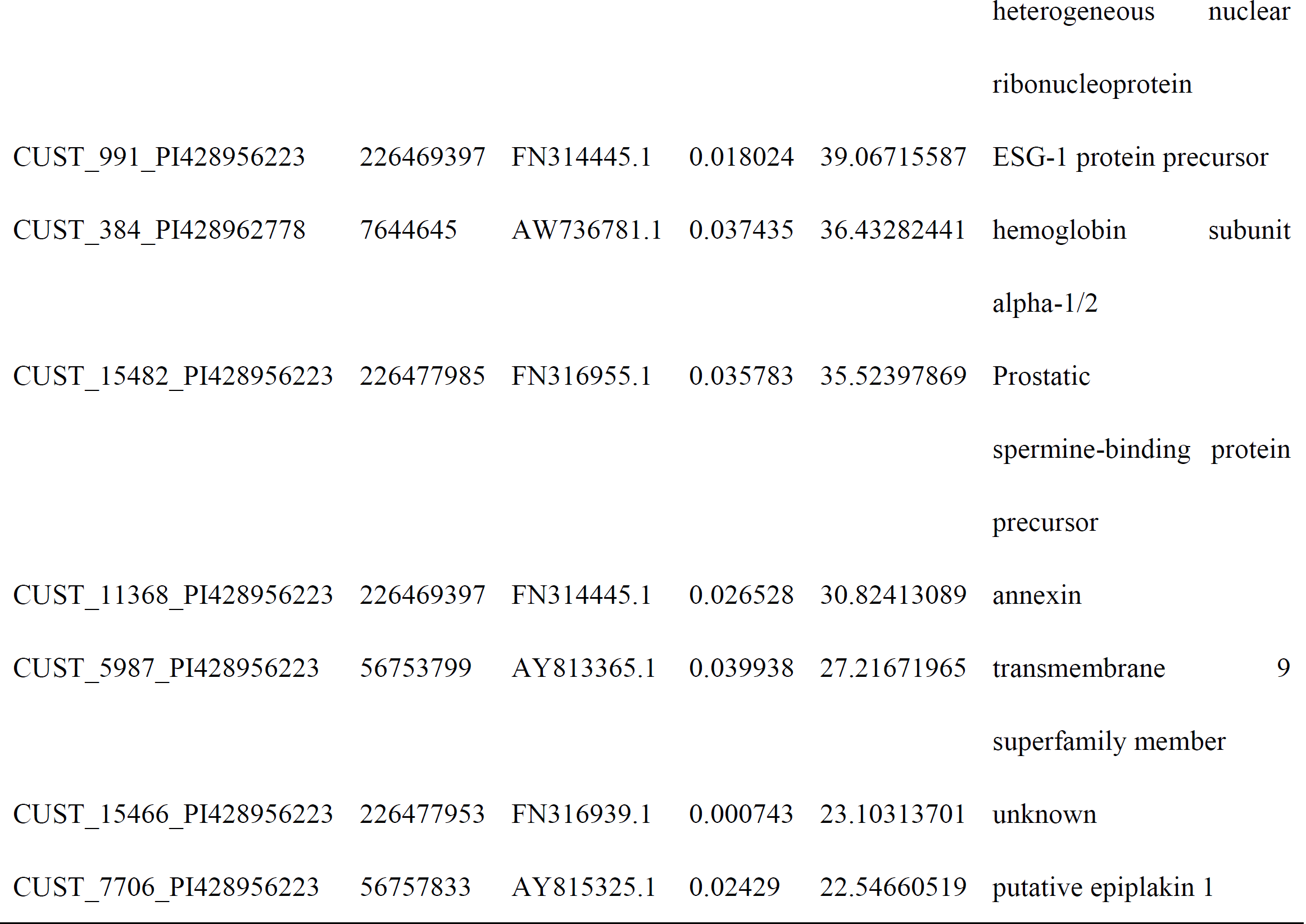
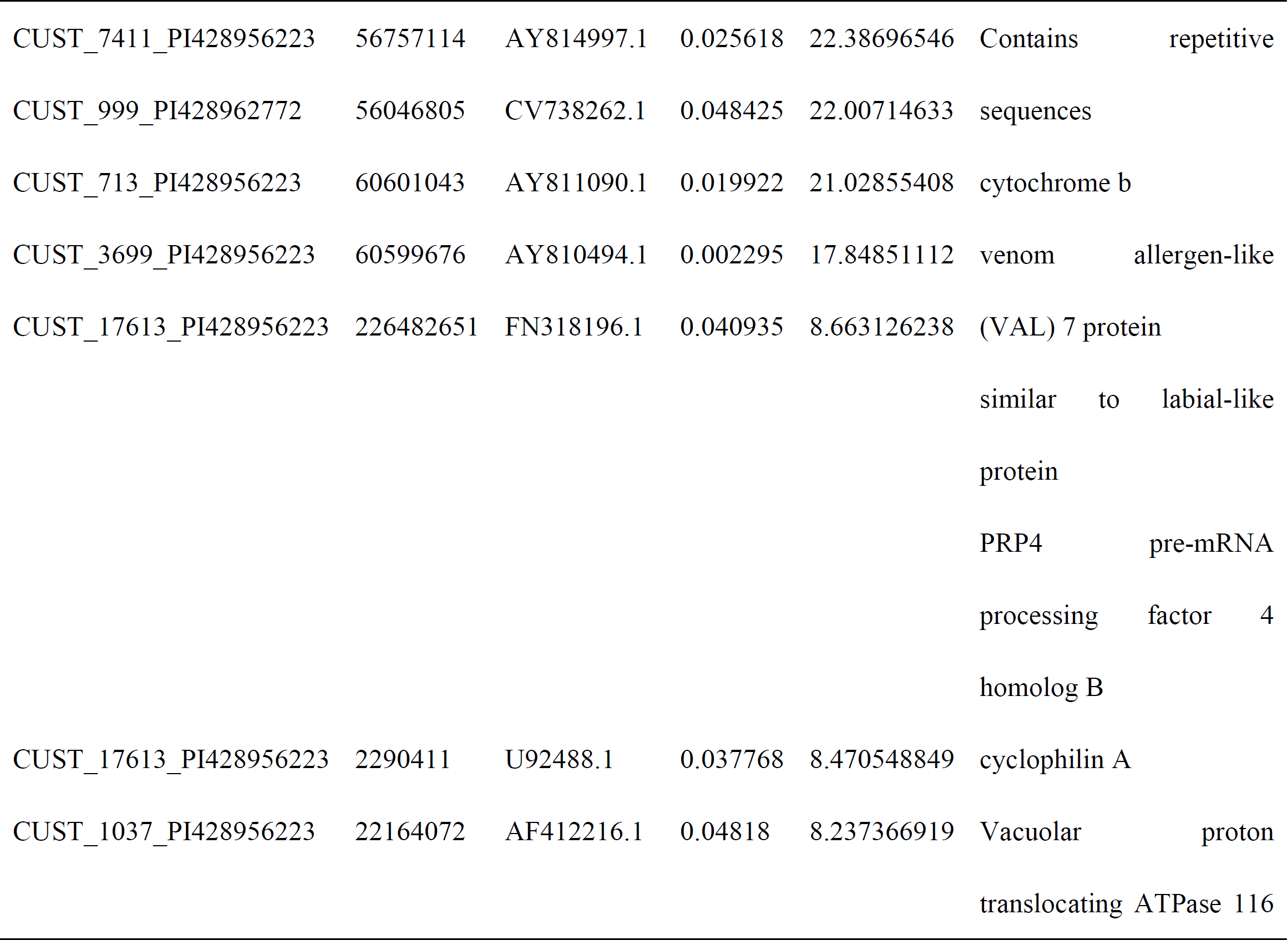
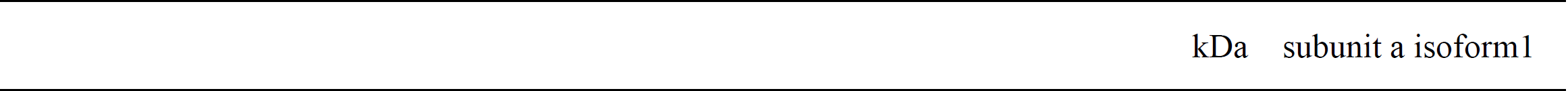
The Top 30 Differentially Expressed Genes in Female S. japonicum after Male-Female Pairing Relative to Levels Prior to Pairing.

### Bioinformatics analysis of differentially expressed genes

To determine the potential function of these upregulated genes, the Gene Ontology (GO) functional categories were assessed. In the biological process GO category, the genes involved in metabolic and biosynthetic processes were more active in females during and after pairing than before pairing, indicating that nutritional acquisition is more crucial for paired than unpaired female trematodes, which is likely a reflection of the oviposition status. In the molecular function GO category, the genes involved in hydrolase activity were more active in females during and after pairing than before pairing. In the cellular component GO category, gene products localized to membrane-bound organelles were more enriched in females after pairing than before or during pairing.

### Confirmation of differentially expressed genes by qPCR analysis

Fifteen differentially expressed genes were selected for confirmation using qPCR. The analysis for each gene was repeated three times, and *PSMD4* was used as the housekeeping gene. After normalization, the relative changes in gene expression were determined using the 2^-ΔΔct^ method. The expression levels of two genes, *annexin* and *tetraspanin-1*, were not consistent between the qPCR and microarray analyses. The results for the remaining thirteen gene expression levels were consistent across qPCR and microarray analyses (Fig 1).

**Fig.1.**
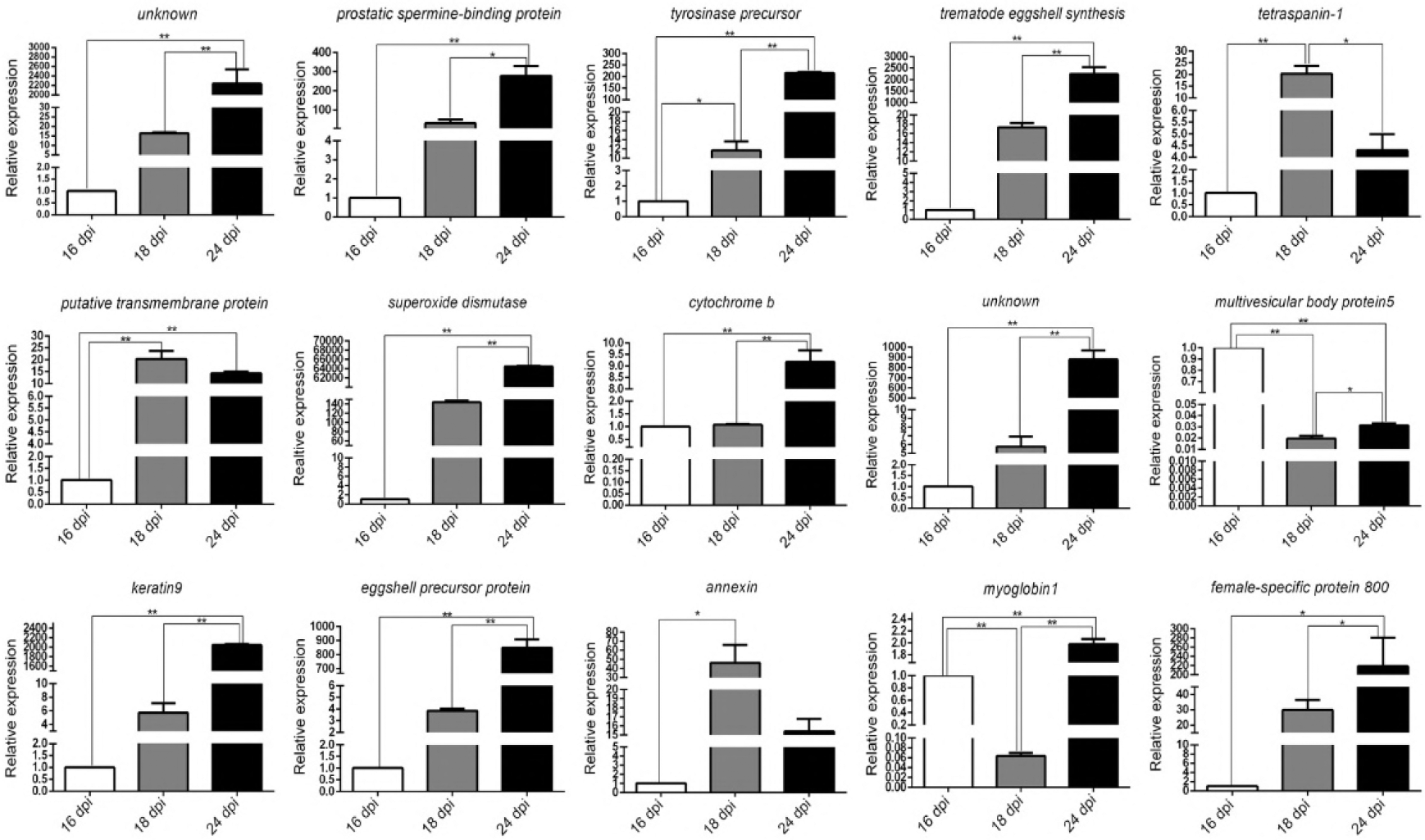
Validation of the Microarray Analysis Results by qPCR. Data are normalized to the internal housekeeping control *PSMD4.* Expression levels of 15 selected genes were determined using the comparative method (2^-ΔΔct^). Data are expressed as the mean ± SEM of three independent experiments (n = 3); independent-sample *t* test, *p < 0.05, *\*\*p* < 0.001.

### Expression of *Sjfs800, cu-zn-SOD, and eggshell precursor protein* in the eggs, cercariae, and schistosomula of females 24 and 42 dpi

The gene expression levels for *Sjfs800*, *cu-zn-SOD*, and *eggshell precursor protein* were determined using qPCR of the eggs, cercariae, schistosomula (at 16 dpi), and female worms 24 and 42 dpi. As above, the results were normalized to the housekeeping gene *PSMD*4, and the relative expression was then determined using the 2^-ΔΔct^ method. We found three genes that were highly expressed in females 42 dpi, and *eggshell precursor protein* and superoxide dismutase were also highly expressed in females 24 dpi. The expression levels of *Sjfs800* were low in the eggs, cercariae, and schistosomula (at 16 dpi) and modestly increased in female worms at 24 dpi, with a further increase at 42 dpi (Fig 2). These results suggested that these genes may be associated with the differentiation and development of *S. japonicum.*

**Fig.2.**
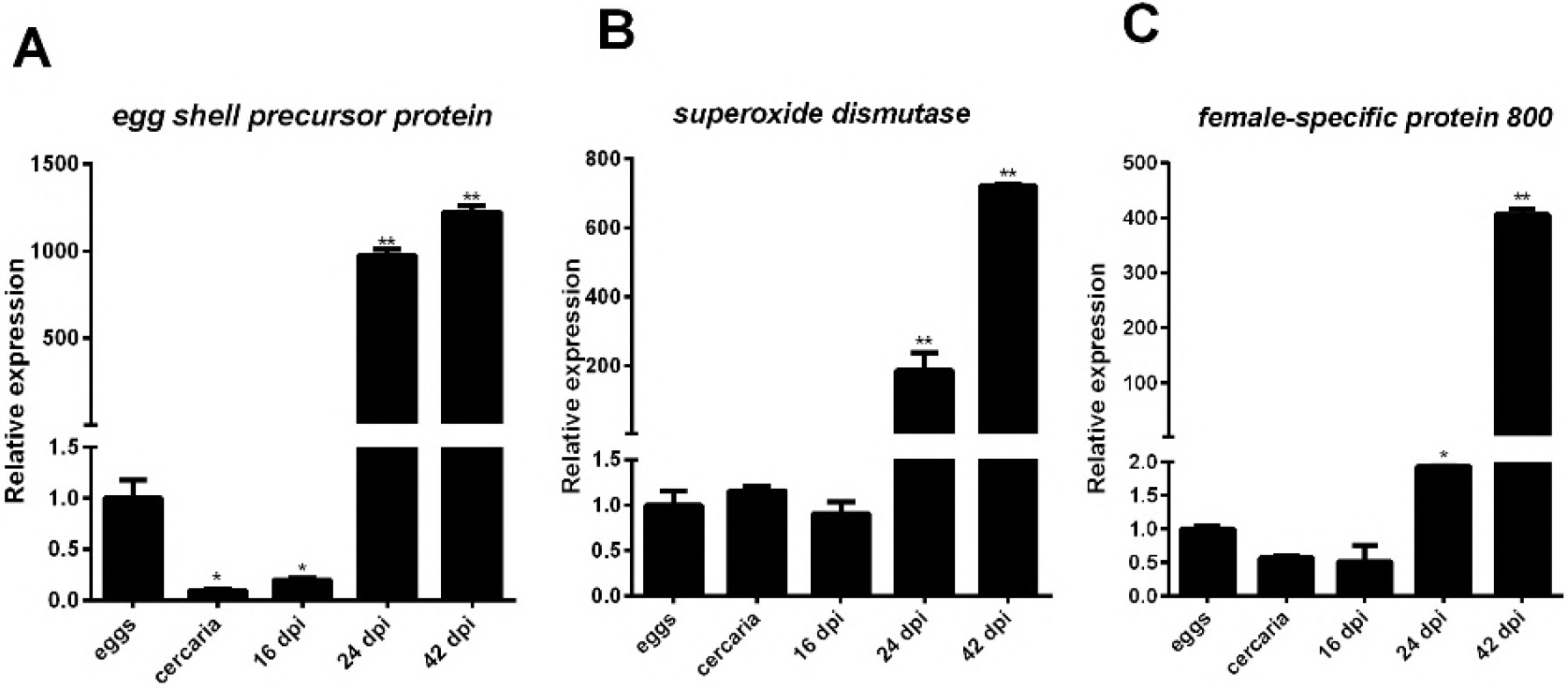
Relative mRNA Expression Levels of Female-Specific Protein 800, Superoxide Dismutase, and Eggshell Precursor Protein at Various Developmental Stages of *S. japonicum* as Measured by qPCR.

The eggs, cercariae, schistosomula (at 16 dpi), and female trematodes 24 and 42 dpi were analyzed (n = 3; *\*p* < 0.05, *\*\*p* < 0.001, compared with the egg group).

### Localization by in situ hybridization of *Sjfs800, cu-zn-SOD*, and *eggshell precursor protein* transcripts in female *S. japonicum*

A situ hybridization analysis showed the transcriptional activities of *cu-zn-SOD, eggshell precursor protein,* and *Sjfs800* in female *S. japonicum* 24 dpi (Fig 3). Although *cu-zn-SOD* mRNA was located throughout much of the whole body, the signal intensity in the ovary and vitellarium was markedly stronger than that in other parts. *Eggshell precursor protein* was substantially expressed in the ovary and ootype and was also found in the vitellarium. The expression of *Sjfs800* mRNA was observed only in the vitellarium.

**Fig 3.**
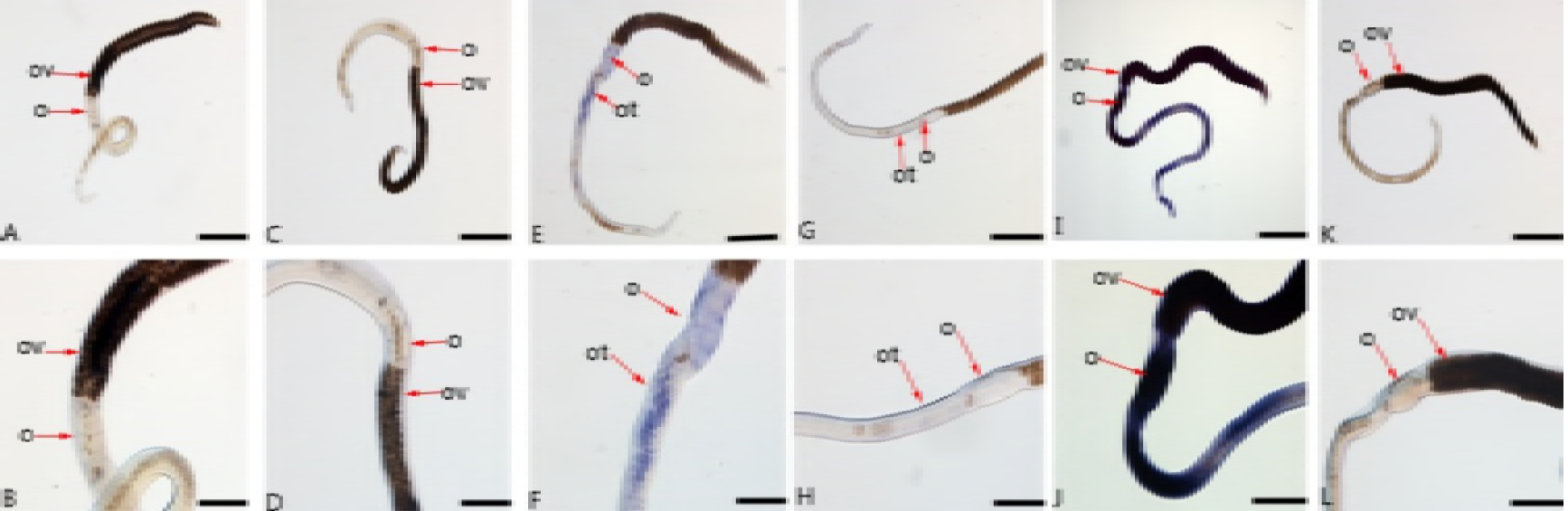
Localization of *Sjfs800, eggshell precursor protein,* and *cu-zn-SOD* Transcripts in Female *S. japonicum* 28 dpi. Images are of whole-mount in situ hybridization results using DIG-labeled antisense (A, B, E, F, I, J) and sense (C, D, G, H, K, L) RNA probes. All three transcripts (A, B, C, D: *Sjfs800;* E, F, G, H: *eggshell precursor protein*; I, J, L, K: *cu-zn-SOD)* were detected. Abundant transcription of *Sjfs800* was observed in the vitellarium; transcription of *eggshell precursor protein* was observed in the ovary and ootype; and abundant transcription of *cu-zn-SOD* was observed in the ovary and the vitellarium. Abbreviations: ov, ovary; vt, vitellarium; ot, ootype. Scale bars: top row, 200 μm; bottom row, 100 μm.

### qPCR analysis of *Sjfs800* mRNA levels after *Sjfs800-specific* siRNA transfection in trematodes 28 dpi

The *Sjfs800* mRNA levels were analyzed by qPCR to determine the effects of the *Sifs800*-specific siRNA transfection. First, the paired 28-dpi worms were transfected with one of the three siRNAs (siRNA1–3) targeting *Sjfs8OO.* After 3 days’ cultivation, *Sjfs8OO* gene transcript levels were determined by qPCR. The reduction in *Sjfs8OO*transcription level following transfection with siRNA1 was 60% of that in the mock transfected group and negative control group, which was the highest efficiency among the three siRNAs tested. Thus, *Sifs800*-specific siRNA1 was used in the ensuing experiments. worms were transfected with *Sifs800*-specific siRNAl or scrambled siRNA in vitro. The results of the qPCR analysis indicated that compared with the scrambled siRNA control group, the *Sifs800*-specific siRNA1-treated group showed an approximately 60% reduction in *Sjfs8OO* mRNA levels on the tenth day later, and this experiment was repeated three times (Fig 4).

### Effects of *Sjfs800* knockdown on pairing rate, egg production, and reproductive organ development

The number of male-female paired worms was counted on the tenth day to determine the effect of knocking down the *Sjfs8OO* gene on the pairing rate. We found that the pairing rate in the *Sifs800*-specific siRNA1 transfected group was significantly lower than that in the scrambled siRNA transfected group. In addition, the number of eggs collected in the medium and counted using light microscopy was reduced approximately 50% in the *Sifs800*-specific siRNAl transfected group compared with that in the scrambled siRNA transfected group (Fig 4).

**Fig.4.**
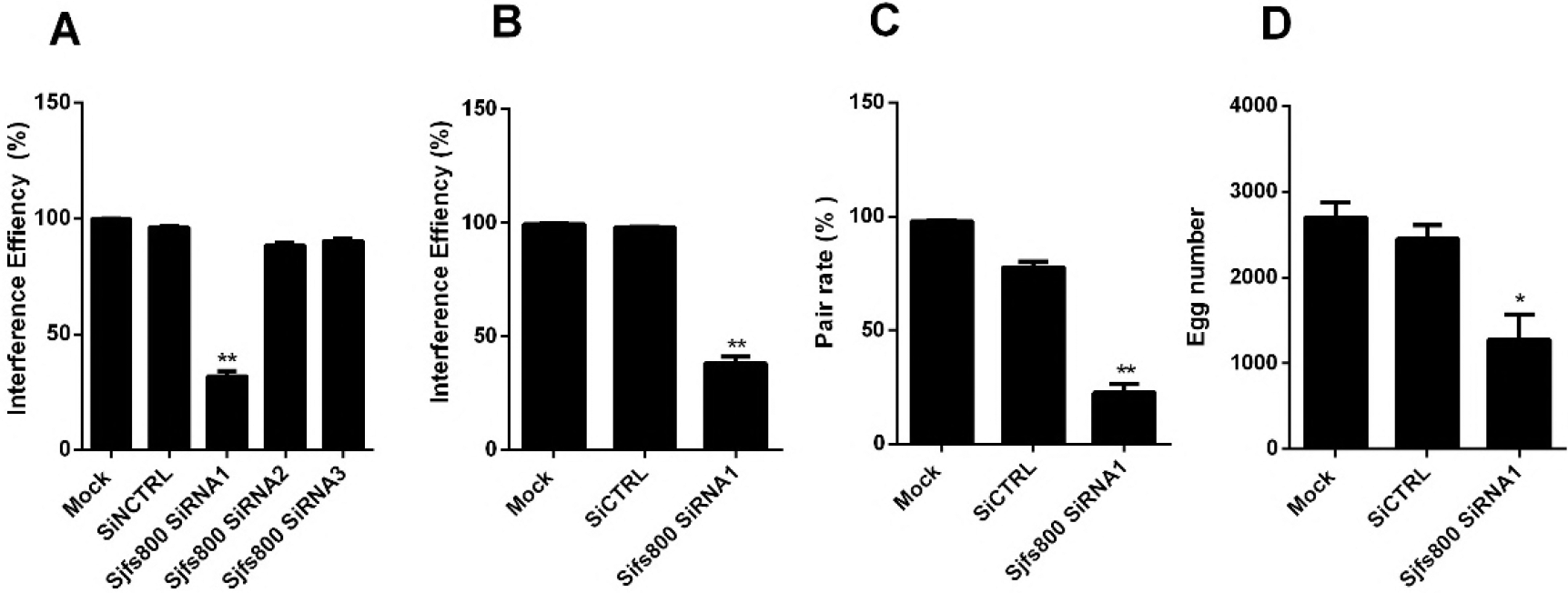
Effects of Sjfs800-specific siRNA Transfection.

A, To determine the Sfs800-specific siRNA with the best interference efficiency in vitro as measured using qPCR, 28-dpi worms were transfected with one of three siRNAs (siRNA1–3) and were harvested 3 days later. The qPCR results showed the effects of siRNA1 were reduced by nearly 60% compared with those in mock-transfected group. B, Effects of siRNA1 on *Sjfs800* mRNA levels in worms were tested 10 days later. *Sjfs800* mRNA levels were normalized to the endogenous control *SjPSMD4.* The qPCR analysis results showed that the *Sjfs800* mRNA levels in the group transfected with *Sjfs800* siRNA1 were reduced by approximately 60% compared with those in the scrambled siRNA-transfected group. C, The male-female pairing rate of worms transfected with *Sjfs800* siRNA1 was reduced by approximately 70% compared with the rate in the scrambled siRN-treated group. D, The eggs in the culture medium were collected on the tenth day and counted using light microscopy. SiNCTRL represents negative control siRNA group; SiCTRL represents scrambled siRNA.

The morphologic changes in the vitellarium of *Sjfs800* siRNA1-treated worms were observed using confocal laser scanning microscopy. The vitellarium was well developed in female worms that were transfected with scrambled siRNA. By contrast, transfection with *Sjfs800* siRNA1 suppressed the development and maturation of the vitellarium, with fewer mature vitelline cells found in the *Sjfs800* siRNA1-treated females than in the controls (Fig 5).

**Fig.5.**
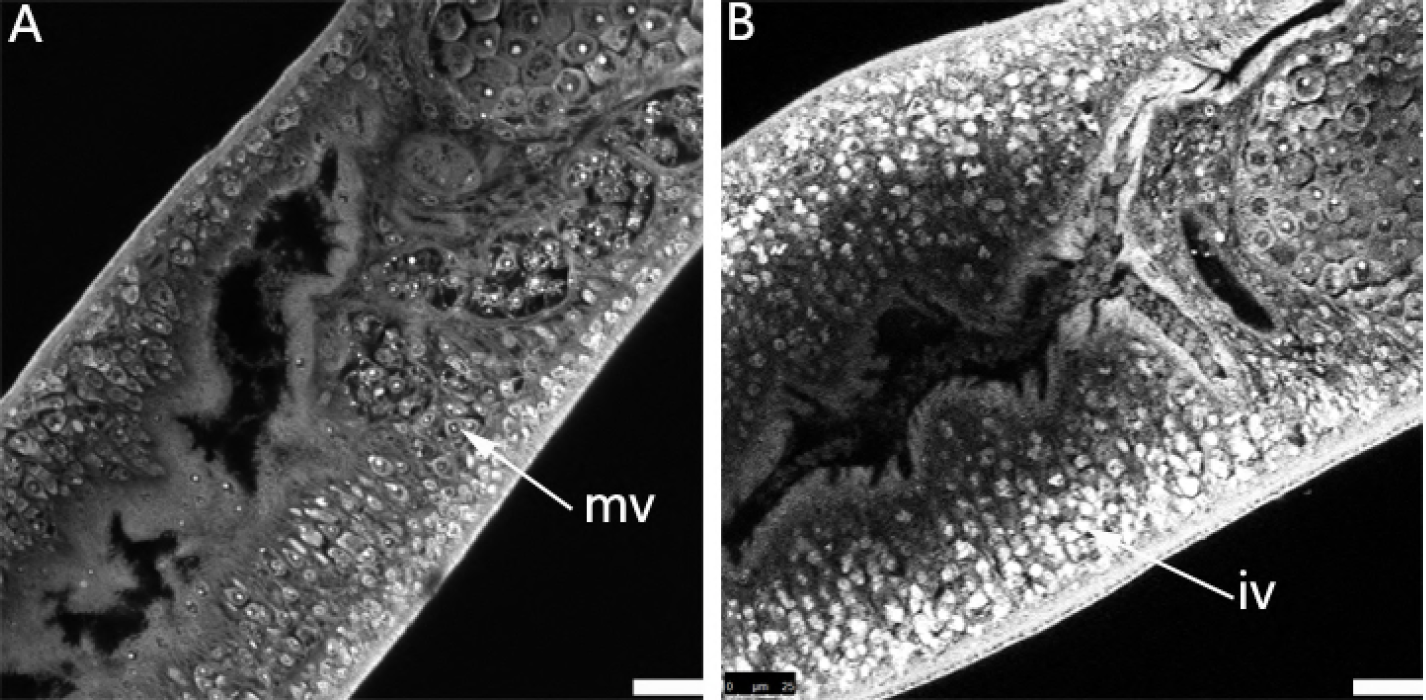
Confocal Scanning Laser Microscopy Images of the Vitellarium in *S. japonicum*. Females After Transfection With *Sjfs800* siRNA1. A, Vitellarium from worms transfected with scrambled siRNA. Arrow points to mature vitelline cells (mv); B, Vitellarium from a trematode transfected with *Sjfs800* siRNA1. Arrow indicates immature vitelline cells (iv). Scale bars, 25 μm.

## Discussion

Paired adult female schistosomes produce a large number of eggs, which are primarily responsible for the schistosomiasis disease pathology and are critical for dissemination of the disease. The reproductive system of female schistosomes has been widely studied at the molecular level, and several studies have described transcriptomes that are differentially expressed before and after pairing in female schistosomes [16, 20-27]. For example, *SmFst* was recently identified as a regulatory molecule in the transforming growth factor β pathway that is pairing dependently transcribed in the male gonad, likely facilitating processes leading to male competence [21]. Lu et al. described pairing-induced processes within the gonads, including stem cell-associated and neural functions, by analyzing gonad-specific and pairing-dependent transcriptomes [22]. In 2017, Wang et al. mapped the dynamic transcriptome changes in male and female *S. japonicum* throughout the sexual developmental process from pairing to maturation to identify biogenic amines and insect-like hormones that regulate reproduction development in *S. japonicum* [16].

In the present study, gene microarray analysis was used to screen for differently expressed genes before and after pairing of the *S. japonicum* female. We found the transcript profiles in 16-dpi females were similar to those at 18 dpi. However, substantial changes in gene expression were observed in 24-dpi females, indicating that virgin female schistosomes undergo marked changes in gene expression before they complete maturation. Genes involved in reproduction, such as those associated with the cell cycle, egg formation, and protein synthesis, were substantially upregulated in the adult female trematodes. This result is consistent with that of others and on the basis of gender-related gene expression patterns reflects that the female, rather than the male, governs egg production [16, 20-27].

The results of our gene microarray analyses showed that the top 30 genes with respect to transcript abundance in 24-dpi females included representatives with proven roles in vitellarium development and egg production, including *eggshell precursor protein*, superoxide dismutase *(SOD),* and *Sjfs800.* The vitellarium, which occupies the posterior two-thirds of the female schistosome body, produces vitellocytes. Vitellocytes supply nutrition to the developing zygote and constituents essential to egg shell construction. Mature vitellocytes join with fertilized oocytes in the ootype, which is where mature eggs are formed [28]. Thus, the development of the vitellarium plays essential roles in the production of schistosome eggs. In the present study, we aimed to determine the role of some of the genes related to vitellarium development in the production of schistosome eggs.

SOD is an antioxidant that removes reactive oxygen species in an organism by converting the superoxide radical to molecular oxygen and hydrogen peroxide. SOD is in the cytoplasm and mitochondria and is also extracellular and is conserved across various species [29]. In the present study, *SOD in* 24-dpi *S. japonicum* females was located primarily in the ovary and vitellarium. Our transcriptome analysis indicated that *SOD* was expressed almost exclusively (>2000-fold enriched) and at high levels in 24-dpi (rather than in 16-dpi or 18-dpi) adult female trematodes, which may be under significant oxidative stress because of vitellarium development and egg production.

The schistosome eggshell is a hardened and tanned structure made from cross-linked proteins. Eggshell formation starts in the ootype. Contractions of the ootype cause the vitelline cells to release their granules, which contain eggshell precursor proteins. The eggshell is shaped by the ootype and strengthened through tyrosinase activity that causes cross-linking of the released eggshell precursor proteins [30]. We found that the *eggshell precursor protein* gene was far more highly expressed in 24-dpi adult female trematodes than in 16-dpi or 18-dpi females. Transcripts of the *eggshell precursor protein* gene were expressed within the ovary and vitellarium, consistent with the findings of a previous study showing similar gene expression [31]. We also observed *eggshell precursor protein* signals in the ootype. Thus, the findings of the present study suggest that eggshell precursor protein maybe play an important role in helping the vitellarium promote eggshell formation.

*Smfs800* gene was first found and identified by Reis et al., who also used in situ hybridization to determine that *Smfs800* mRNA was expressed only in female vitelline cells [32], suggesting that Smfs800 may play role in egg development. However, little is known about the functions of fs800 in vitellarium development and egg production. In the present study, *Sjfs800* mRNA was located in mature vitelline cells. Some developmental defects, especially a reduced number of mature vitelline cells in the vitellarium, were observed after *Sjfs800* gene knockdown by siRNA, and the pairing rate and oviposition rate were also significantly decreased. Therefore, we concluded that fs800 is vital for vitelline cell development and maturation and that maturation of the vitellarium is required for *S. japonicum* females to produce eggs. The number male-female pairings was reduced by approximately 70% after *Sjfs800* siRNA transfection. The results of some studies have indicated that *Sjfs800* may be a molecule downstream of *S. japonicum* Nanos1 or Abl tyrosine kinase activity [33-35].

In conclusion, our study showed substantial differences in the expression levels of some genes in the *S. japonicum* female before and after male-female pairing, including genes related to vitellarium development, which can affect pairing, sexual maturation, and egg production. These results provide a deeper understanding of the reproductive biology of schistosomes and may lead to the development of novel approaches for the prevention and treatment of schistosomiasis.

